# Modeling biases from low-pass genome sequencing to enable accurate population genetic inferences

**DOI:** 10.1101/2024.07.19.604366

**Authors:** Emanuel M. Fonseca, Linh N. Tran, Hannah Mendoza, Ryan N. Gutenkunst

## Abstract

Low-pass genome sequencing is cost-effective and enables analysis of large cohorts. However, it introduces biases by reducing heterozygous genotypes and low-frequency alleles, impacting subsequent analyses such as demographic history inference. We developed a probabilistic model of low-pass biases from the Genome Analysis Toolkit (GATK) multi-sample calling pipeline, and we implemented it in the population genomic inference software dadi. We evaluated the model using simulated low-pass datasets and found that it alleviated low-pass biases in inferred demographic parameters. We further validated the model by downsampling 1000 Genomes Project data, demonstrating its effectiveness on real data. Our model is widely applicable and substantially improves model-based inferences from low-pass population genomic data.

## Introduction

Enabled by reduced sequencing costs, population genetics has experienced a revolution, from focusing on a limited number of loci to now encompassing entire genomes (Maddison et al. 1992; Reid et al. 2016; Marchi et al. 2022). Yet researchers must still trade off a) the extent of the genome to be sequenced, b) the depth of coverage for each sample, and c) the number of sequenced samples (Lou et al. 2021; Martin et al. 2021; Duckett et al. 2023). One way to address this trade off is to sequence one reference sample at high coverage depth while sequencing others at lower depth (Lou et al. 2021). Low-pass sequencing, in which the genome is sequenced at a lower depth of coverage, avoids many of the financial, methodological, and computational challenges of high-pass sequencing (Li et al. 2011). Furthermore, limited availability of DNA can also make high depth impractical, especially for ancient samples and museum or herbarium specimens (Mota et al. 2023).

Despite its advantages, low-pass sequencing may lead to an incomplete and biased representation of genetic diversity within a population (e.g., Vieira et al. 2013; Fox et al. 2019). Low-frequency genomic variants may not be detected (Fumagalli 2013), and genotypes may be less accurate (Nielsen et al. 2011). Low-pass sequencing increases the likelihood of miscalling heterozygous loci as homozygous (Duitama et al. 2011; Gorjanc et al. 2015), due to a lack of sufficient reads on homologous chromosomes to distinguish between different alleles at a given locus. These issues can then bias downstream analyses. It is thus important for analysis methods to accommodate low-pass sequencing (see Carstens et al. 2022 for a discussion of related issues).

The allele frequency spectrum (AFS) is a powerful summary of population genomic data (Sawyer & Hartl 1992; Wakeley 2009). Briefly, the AFS is matrix which records the number alleles observed at given frequencies in a sample of individuals from one or more populations. The AFS is often the basis for inferring demographic history (Gutenkunst et al. 2009) or distributions of fitness effects (Kim et al. 2017). In low-pass sequencing, the loss of alleles and the excess of homozygosity can bias the estimation of the AFS (Fumagalli 2013) and thus those inferences.

To address the challenges of low-pass data, several tools have emerged (Bryc et al. 2013; Blischak et al. 2018; Meisner & Albrechtsen 2018), with one of the most widely adopted being ANGSD (Korneliussen et al. 2014). ANGSD offers a diverse range of analyes tailored for low-pass sequencing data. To infer an AFS, ANGSD uses sample allele frequency likelihoods, which can be computed either directly from raw data or, more frequently, from genotype likelihoods (Nielsen et al. 2012). These likelihoods quantify the probability of observing the complete set of read data for multiple individuals at specific genomic sites, given particular sample allele frequencies (Nielsen et al. 2012; Korneliussen et al. 2014), enabling ANGSD to estimate allele frequencies. While ANGSD has proven its utility, limitations exist. For example, many analyses rely on distinguishing different types of variant sites (such a synonymous versus nonsymonyous) which the developers of ANGSD recommend against. Moreover, in some cases unbiased estimation of the AFS may be difficult or impossible.

Rather than attempting to estimate an unbiased AFS from low-pass data, we developed a probabilistic model of low-pass AFS biases and incorporated it into the population genomic inference software dadi (Gutenkunst et al. 2009). Our model is based on the multi-sample genotype calling pipeline of the Genome Analysis Toolkit (GATK), the most widely used tool for calling variants from read data (McKenna et al. 2010; Auwera & O’Connor 2020). We assessed the accuracy of our model using simulated low-depth data as well as subsampled data from the 1000 Genomes Project (Fairley et al. 2020, https://www.internationalgenome.org/). We found that our model accurately captures low-depth biases in the AFS and enables accurate inference of demographic history from low-pass data.

### Model for Low-pass Biases

When biases arises from low-pass sequencing, the AFS may be affected by both the loss of low-frequency variants and the misidentification of heterozygous individuals as homozygous. These two effects result in a deficit of variant sites and misleading shifts in allele frequencies, respectively. Moreover, the data must often be subsampled to generate an AFS for analysis, because not all individuals will be called at all sites. We account for these distortions by sequentially modeling the probabilities of a variable site being called, of that site having enough called individuals for subsampling, and of having its allele frequency misestimated. The specific choices in our model are motivated by the default GATK multi-sample calling algorithm, in which information from all samples is used to identify whether a site is variant. In particular, we assume that a site will only be called as variant if at least two alternate allele reads are observed. Once a site is identified as variant, an individual will be called as missing if zero reads are observed, homozygous if all reads correspond to a single allele, and heterozygous if at least one reference and one alternate read are observed. For simplicity, we first describe the case of sequencing *n*_*seq*_ individuals from a single population.

Consider a site in which the true alternate allele count within our sample of *n*_*seq*_ individuals is *f*. Those *f* alternate alleles can be distributed among the 2*n*_*seq*_ sampled alleles in many ways. To quantify those ways, we define the partition function ℙ_*nseq*_(*f*), which is an array of integer partitions with *n* entries that sum to the allele frequency *f* such that all entries in the partition are 0, 1, or 2 (corresponding to the possible genotype values). For example, the partitions defined by ℙ_4_(3) are [2, 1, 0, 0] and [1, 1, 1, 0]. Each possible partition within ℙ_*nseq*_(*f*) can occur in 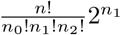 ways, where *n*_0_, *n*_1_, and *n*_2_ denote the number of partition entries equal to 0, 1, or 2. (The factor of 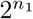 accounts for the two possible haplotypes the alternate allele could lie on in each heterozygote.) The corresponding probability of each partition within ℙ_*nseq*_(*f*) is then the number of ways it can occur divided by the total over all partitions within ℙ_*nseq*_(*f*).

Let 𝔻 denote the distribution of read depth *d* within the population sample, which we assume to be shared among all individuals. For an individual homozygous for the alternate allele, the probability of observing *a* alternate reads is simply 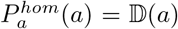. For a heterozygous individual, the probability of zero alternate reads is

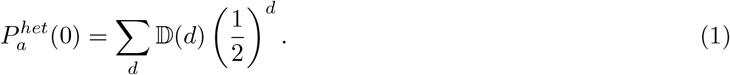

Here we sum over the distribution of depths, and at each depth each read has a 1*/*2 chance of containing the reference allele, so the probability of all reads being reference is (1*/*2)^*d*^. Similarly, the probability of exactly one alternate read is

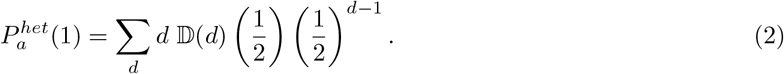

Note that for depth *d*, there are *d* possible configurations with one alternate read and *d* − 1 reference reads.

For a given partition within ℙ_*nseq*_(*f*) that has true genotype counts *n*_0_, *n*_1_, and *n*_2_, there are multiple ways of failing to identify the variant site. The probability of zero reads supporting the alternate allele is

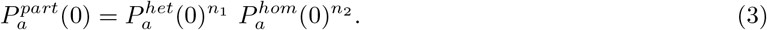

The probability of exactly one read supporting the alternate allele is

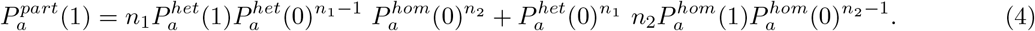

Here the two terms account for the probability that the alternate read occurs in one of the heterozygotes or homozygotes, respectively. The overall probability of not calling a variant site for a given partition is thus 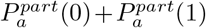. And the overall probability of not calling a variant site with a given true allele frequency *f* is the sum of these probabilities over partitions ℙ_*nseq*_(*f*), weighted by the partition probabilities. For any given coverage distribution, the probability of calling a variant site increases rapidly with allele frequency *f* (Fig. S1).

When analyzing low-pass data, generating an AFS for the full sample size *n*_*seq*_ may result in the loss of many sites where not all individuals were called. Consequently, it is common to subsample the data to some lower sample size *n*_*sub*_; only sites with calls for at least *n*_*sub*_ individuals can then be analyzed. The probability a site can be analyzed is independent of the allele frequency and is

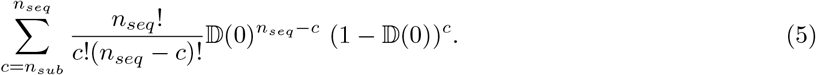

Here we sum the probability that exactly *c* individuals have at least one read at this site over all potential values of the number of covered individuals *n*_*sub*_ ≤*c* ≤*n*_*seq*_. From this point onward, we consider partitions ℙ_*nsub*_(*f*) over the subsampled individuals.

Once a site as called as variant, low-pass sequencing can bias the estimation of the allele frequency at that site, if one or more heterozygotes are miscalled because all their reads are reference or alternate. For each heterozygous individual, this occurs with total probability

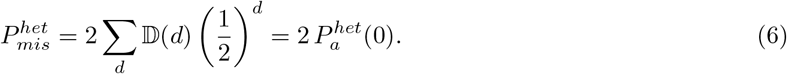

For a partition with *n*_1_ true heterozygotes, the number of miscalled heterozygotes 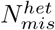 is binomially distributed with mean 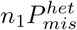. Each miscalled heterozygote has equal chance of being called as homozygous reference or alternate, so the number of miscalls to homozygous reference 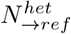 is binomially distributed with mean 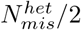, and the number of miscalls to homozygous alternate is 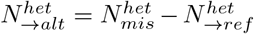. The net change in estimated alternative allele frequency is then 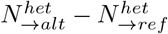.

The biases caused by low-pass sequencing do not depend on the underlying AFS; for each true allele frequency a given fraction will always, on average, be miscalled as any given other allele frequency. The correction above can be thus be calculated once for a given data set then applied to all model AFS generated, for example, during demographic parameter optimization. For efficiency, we calculate and cache an *n*_*seq*_ by *n*_*nub*_ transition matrix that can be multiplied by any given model AFS for *n*_*seq*_ individuals to apply the low coverage correction. When analyzing multiple populations, we calculate and apply transitions matrices for each population, because variant calling is independent among populations once a variant has been identified. Variant identification is, however, not independent among populations, which we address using simulated calling described next.

When calculating the probability of miscalling a heterozygote (Eq. 6), the correct distribution of depth is not simply 𝔻(*d*); rather it is the distribution conditional on the site being identified as variant. The lower the true allele frequency, the more these distributions will differ. The conditional distribution is complex to calculate, particularly when multiple populations are involved. Instead, for true allele frequencies for which the probability of not identifying is above a user-defined threshold (by default 10^−2^), we simulate the calling process rather than using our analytic results. For multiple populations, we calculate this threshold assuming that a variant must be identified independently in all populations, which gives a lower bound on the true probability of not identifying. To simulate calling, for a given true allele frequency (or combination in the multi-population case) we simulate reads (default 1000) using the coverage distribution 𝔻(*d*) and simulate variant identification and genotype calling for each potential partition of genotypes across the populations, proportional to its probability. For each combination of input true allele frequencies simulated, we estimate and store probability of each potential output allele frequency. These distortions are then applied in place of the transition matrices from the analytic model.

For inbred populations, there is an excess of homozygotes compared to the Hardy-Weinberg expectation, which reduces biases associated with low-pass sequencing. In this case, we follow Blischak et al. (2020) and within each genotype partition calculate the probability of reference homozygotes, heterozygotes, and alternate homozygotes using results from Balding & Nichols (1995, 1997), given the inbreeding coeffficient *F*. The partition probability is then multinomial given these probabilities. In these calculations, we approximate the population allele frequency by the true sample allele frequency. Because calculation of the low-pass correction is expensive compared to typical normal model AFS calculation, we pre-calculate and cache transition matrices and calling simulations. But inbreeding is often an inferred model parameter, to be optimized during analysis. In this case, users can specify an assumed inbreeding parameter for the low-pass model, optimize the inbreeding parameter in their demographic model, update the inbreeding coefficient assumed in the low-pass model, and iterate until convergence.

## Results

### Low-pass sequencing biases the AFS

We used simulated data to assess the biases introduced by low-pass sequencing with GATK multi-sample calling, along with our model of those biases. For a simulated population undergoing growth (Fig. S2A), low-pass sequencing reduces the number of observed low-frequency alleles (Fig. 1). Our model accurately captures these biases (Fig. 1). In contrast with our model, ANGSD attempts to reconstruct the true AFS from low-pass data. In our simulations, ANGSD reconstructed the mean shape of the AFS well, but it introduced dramatic fluctuations into the reconstructed AFS at low depth (Fig. 2).

**Figure 1:**
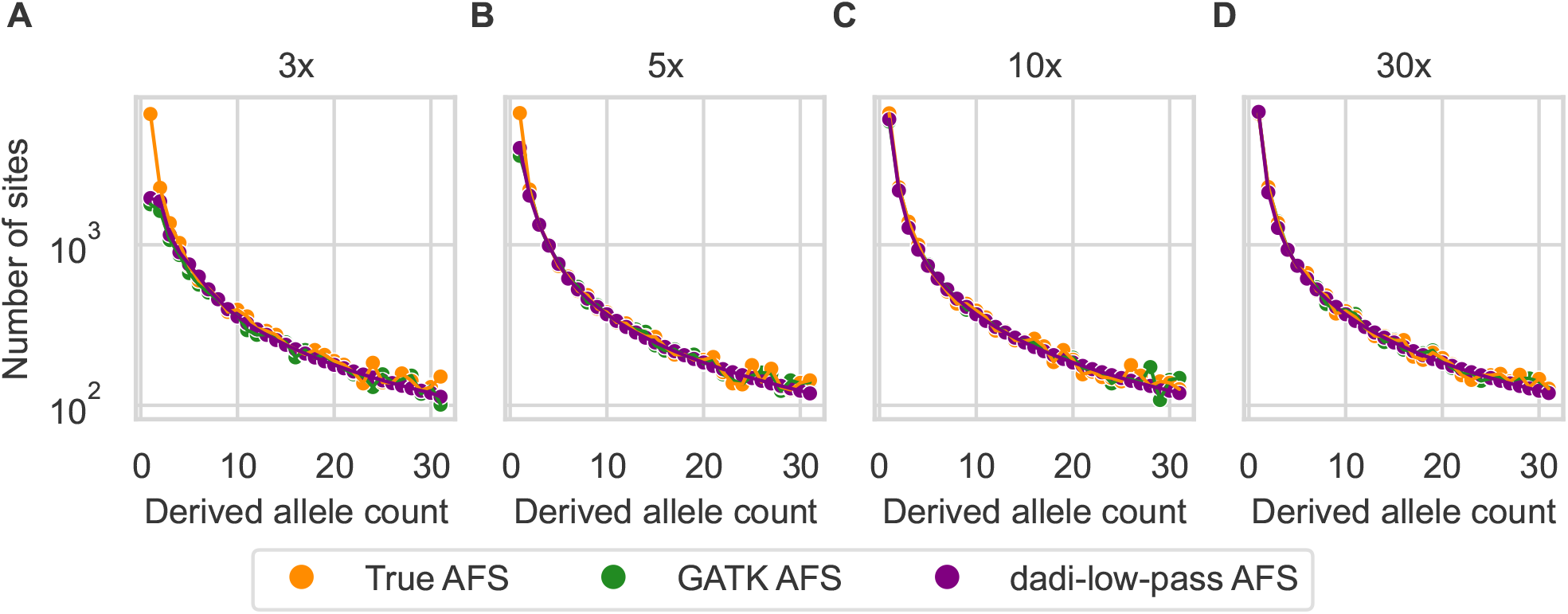
The low-pass AFS is biased, which our model captures. Simulated sequence data from an exponential growth demographic model for 20 individuals were called by GATK and subsampled to 16 individuals (to accommodate missing data at low depth). The GATK-called AFS (green) is biased compared to the true AFS (orange), and our dadi model for low-pass sequencing (purple) fits those biases well. Coverage was (A) 3×, (B) 5×, (C) 10×, and (D) 30×.

**Figure 2:**
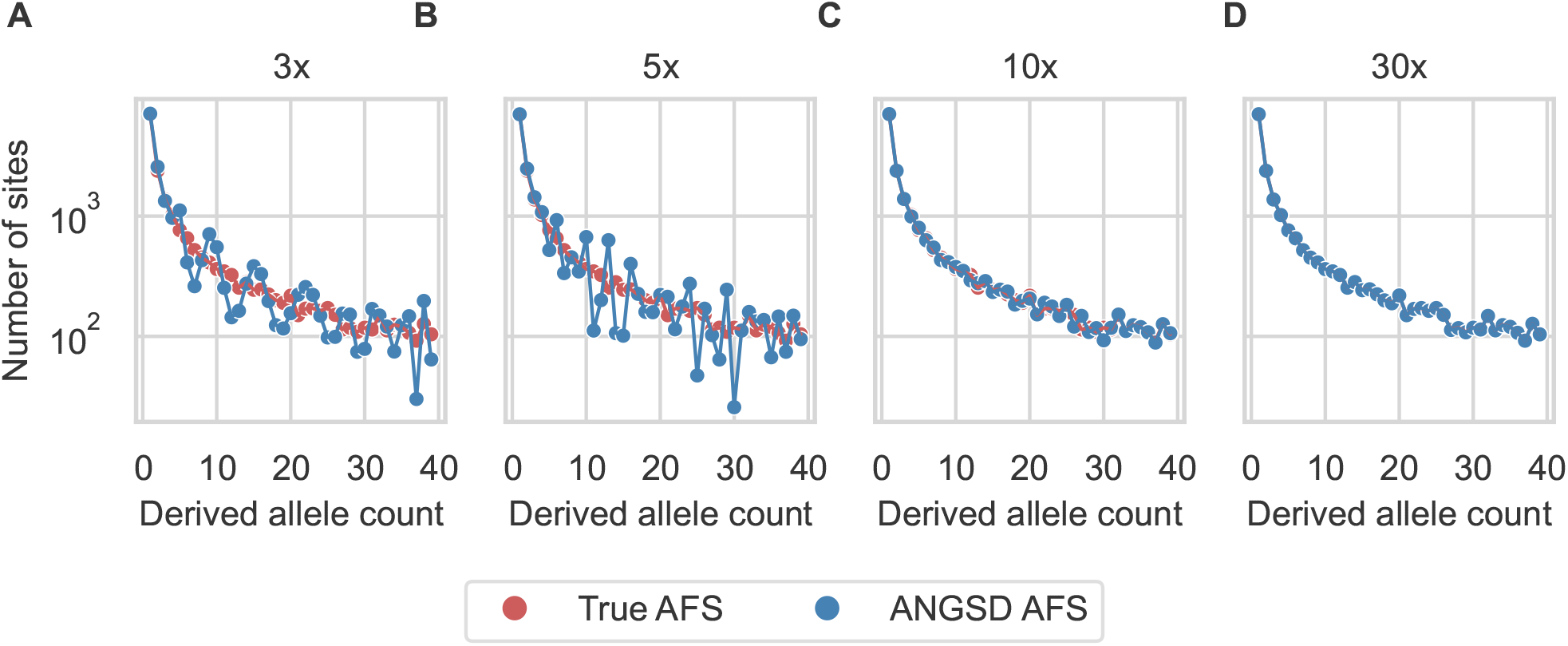
ANGSD corrects for low-pass bias of the AFS, but introduces fluctuations. For the same simulations as Fig. 1, ANGSD (blue) was used to reconstruct the true AFS (red). Coverage was (A) 3×, (B) 5×, (C) 10×, and (D) 30×.

When a pair of populations undergoing a split and isolation (Fig. S2B) is analyzed through a joint AFS, similar low coverage biases occur (Fig. S3). Again, our model corrects those biases well (Fig. S3). Similar to the single-population case, ANGSD also introduces large fluctuations in the joint AFS S4).

Low-pass biases are expected to be smaller in inbred populations, due to the reduction of heterozygosity. In a simulated population recovering from a bottleneck with inbreeding (Fig. S2C), biases are still observed, which our model corrects (Fig. S5). Again, ANGSD introduced large fluctuations in low-pass AFS, beyond those expected from inbreeding (Fig. S6).

### Demographic history inference from low-pass AFS

To assess effects on inference, we first fit demographic models to single-population data simulated under the same growth model as our prior simulations (Fig. S2A). When not modeling low-pass biases, the final population size was underestimated (Fig. 3A), consistent with a deficit of low-frequency alleles. The timing of growth onset was also inaccurately inferred, underestimated at 3× depth and overestimated at 5× depth (Fig. 3B). When the same data were fit with our low-pass model, both model parameters were accurately recovered (Fig. 3A&B) even at the lowest depth. Fits to the AFS reconstructed by ANGSD also yielded accurate model parameters (Fig. 3A&B).

**Figure 3:**
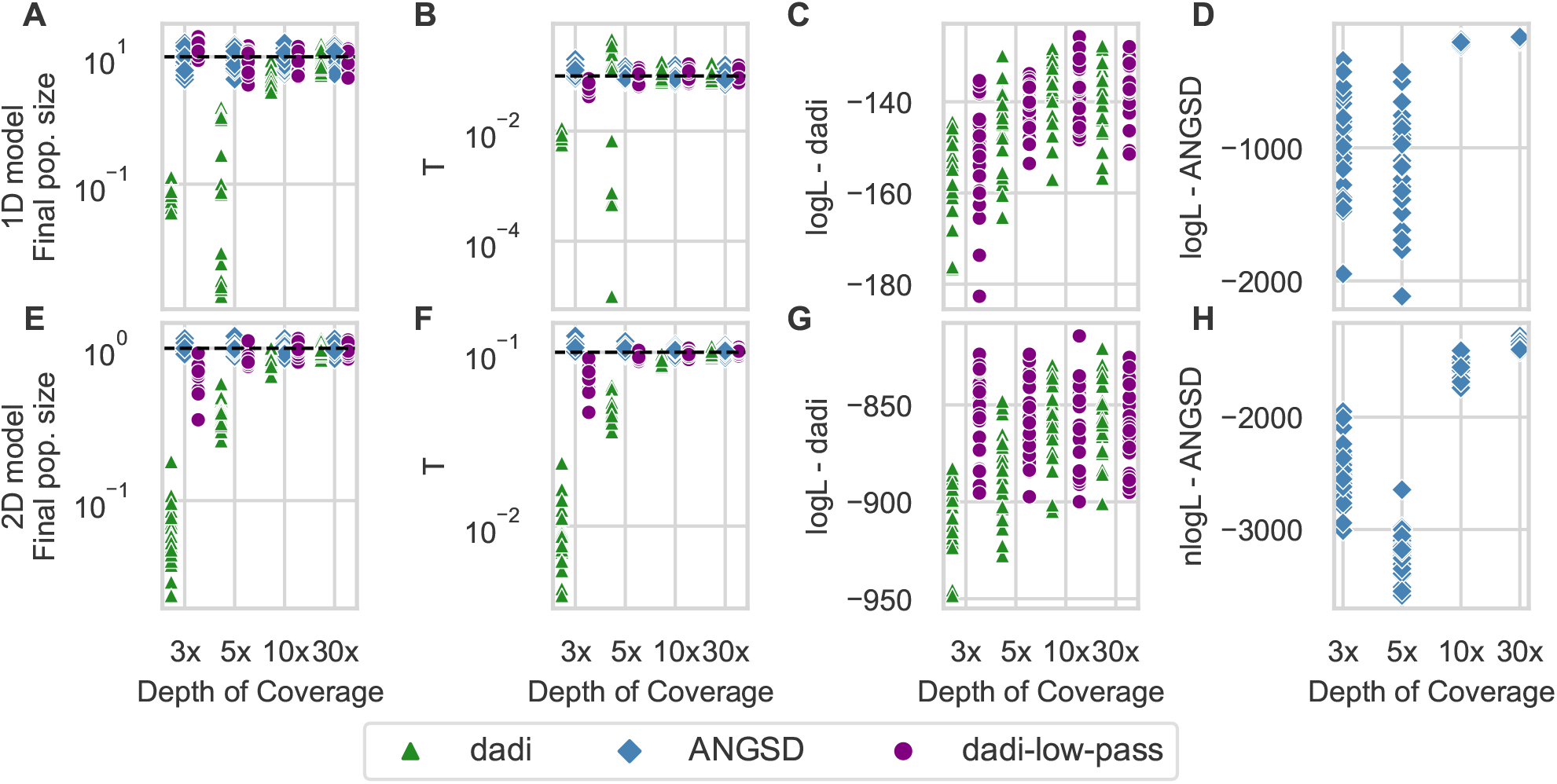
Our low-pass model and ANGSD enable accurate demographic parameter inference. A&B: From data simulated under a single-population growth model, the final population size and time of growth onset (*T*) were accurately inferred using our low-pass model and a GATK-called AFS or using normal dadi and an ANGSD-called AFS. But they were biased if low depth was not accounted for when fitting a GATK-called AFS. (Dashed horizontal lines are simulated true values.) C: The likelihoods using the GATK AFS were similar whether or not low-pass biases were modeled. D: The fluctuations introduced into the AFS by ANGSD caused low likelihoods at low depth. E-H: For a two-population split with isolation model, similar results were found, although inferences from our low-pass model were slightly biased at 3× coverage.

The logarithm of the likelihood is commonly used to assess the quality of model fit. ANGSD reconstructs the AFS for the full sequenced sample size, while we subsample in our approach to deal with missing genotypes, so the likelihoods are not directly comparable. The likelihoods of models fit to the subsampled GATK data were similar whether or not low-pass biases were modeled (Fig. 3C), suggesting that the likelihood itself cannot be used to detect unmodeled low-pass bias. When fitting AFS estimated by ANGSD, likelihoods were much lower at low coverage than high coverage (Fig. 3D), likely driven by the fluctuations ANGSD introduced into the estimated AFS (Fig. 2).

For two-population data simulated under an isolation model (Fig. S2B), similar results were found. Fitting the observed low-pass AFS with our model enabled accurate parameter inference (although there was some bias at 3× coverage) as did fitting the AFS estimated by ANGS (Fig. 3D). As in the single-population case, likelihoods were substantially lower when fitting the ANGSD-estimated AFS, consistent with introduced fluctuations in the AFS (Fig. S4).

For one-population data simulated under a growth model with inbreeding (Fig. S2C), failing to correct for low-pass biases at low inbreeding (*F* = 0.1 or *F* = 0.5) led to similar biases as with no inbreeding, which our low-pass model corrected (Fig. S7). For high inbreeding (*F* = 0.9), the impact of low-pass sequencing on accuracy was smaller, because inbreeding reduces heterozygosity (Fig. S7).

When applying our low-pass bias correction, the user must specify a value for inbreeding, while they may separately estimate it during demographic parameter optimization. We tested the impact of misspecifying inbreeding in the low-bias correction using data simulated with moderate inbreeding of *F* = 0.5. Large inbreeding values were inferred if inbreeding was initially underestimated in the low-coverage model, and small values were inferred if inbreeding was initially overestimated (Fig. S8C). A substantial difference between the inbreeding coefficient used for correction and the inferred value thus suggests that the assumed inbreeding coefficient was not optimal. Users can thus iterate and update the value assumed in the low-pass correction to converge to a best inference of inbreeding.

### Analysis of human data

To empirically validate our approach and compare with ANGSD, we used chromosome 20 sequencing data from the 1000 Genomes Project, focusing on two sets of samples: Yoruba from Ibadan, Nigeria (YRI) and Utah residents of Northern and Western European ancestry (CEU). We inferred a single-population two-epoch demographic model (Fig. S9A) from the YRI samples, and a two-population isolation-with-migration model (Fig. S9B) from the combined YRI and CEU samples. To mimic low-pass sequencing, we subsampled the original high-depth data (which averaged 30× per site per individual) to create data with low to medium depth.

As with simulated data, the observed AFS from low-pass subsampled data was biased compared to high-pass data (Fig. 4A). Using the GATK pipeline, low-pass data yielded few low- and high-frequency derived alleles. In contrast to the simulated data, on these real data ANGSD failed to recover the correct number of low-frequency alleles at 3× and 5× depth, while still introducing large fluctuations at intermediate frequencies (Fig. 4B).

**Figure 4:**
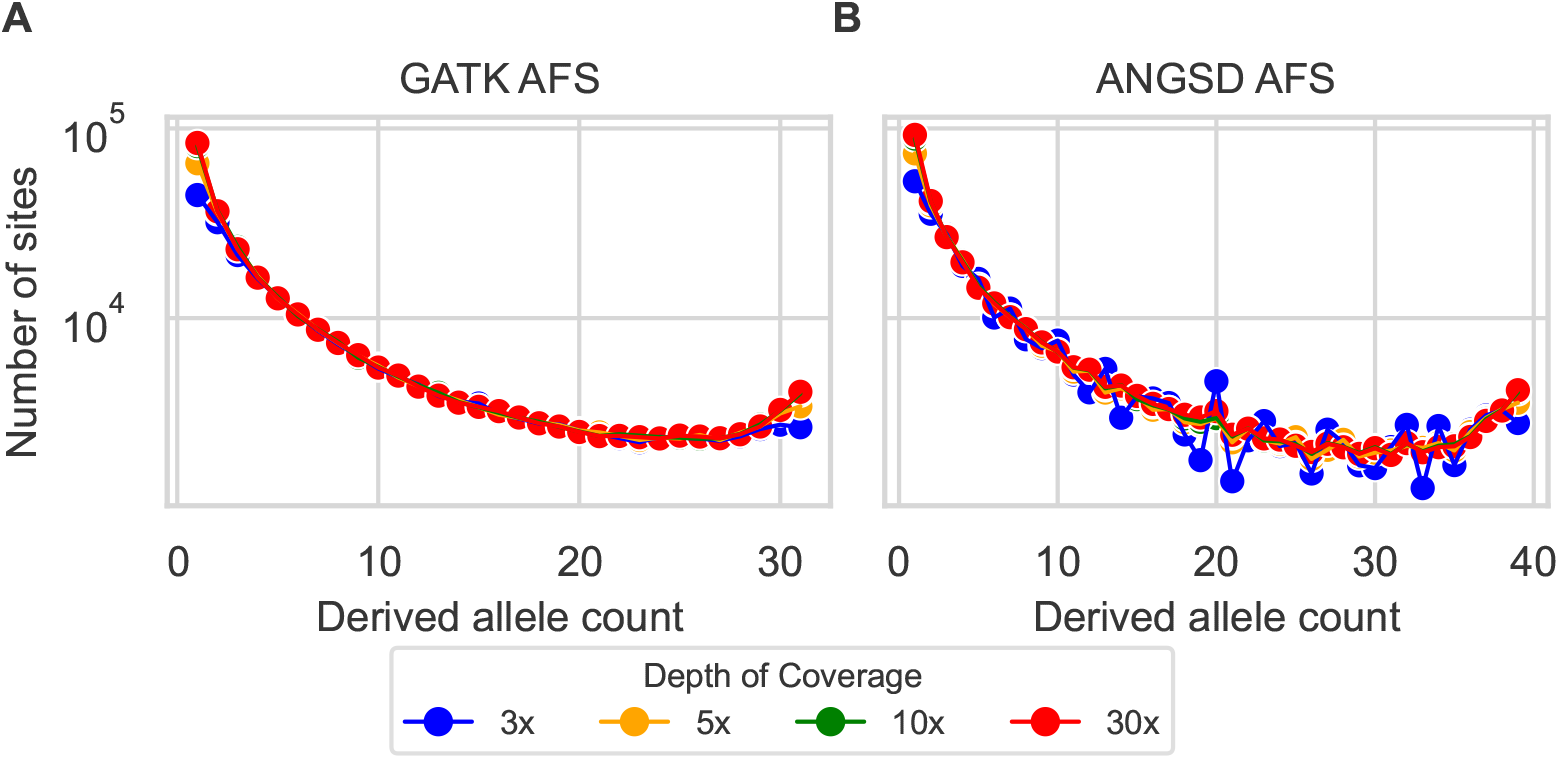
Allele frequency spectra from 20 YRI samples versus subsampled sequencing depth. A: Spectra generated using the GATK pipeline and subsampled to 32 haplotypes to accomodate missing genotypes. B: Spectra generated using ANGSD genotype likelihood optimization with BAM files input.

If low-pass biases were corrected for, we expected the inferred demographic parameters from subsampled low-pass data to match those from the original high-pass data. For the two-epoch model fit to YRI data, we found that with a GATK-called AFS and no low-pass model (Table 1), the inferred population sizes were biased downward and the times were inaccurate, similar to the growth model fit to simulated data. With the low-pass model, inferred values for low depth were similar to those for high depth, with some deviation at 3× (Table 1). Results from fitting ANGSD-estimated spectra were similar to not modeling low depth, suggesting that ANGSD is ineffective for these data (Table 1). As with simulated data, the likelihoods for ANGSD at low depth were also low.

**Table 1:**
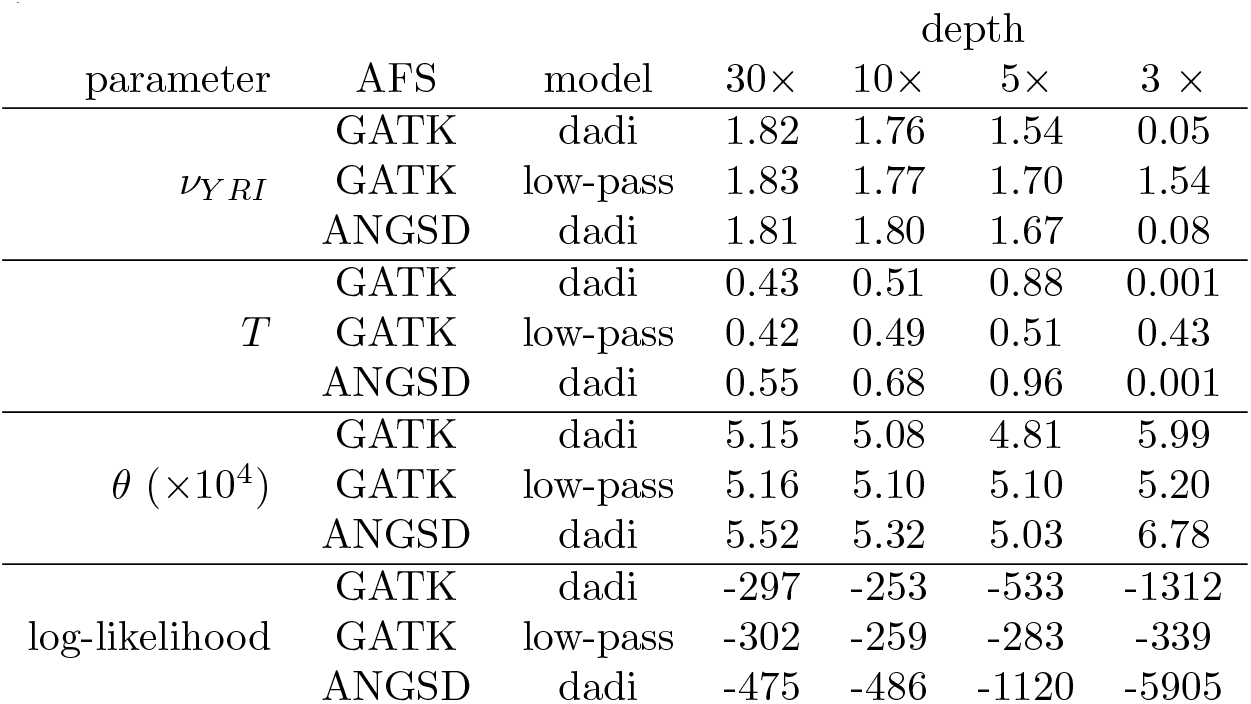
One-population YRI model analysis results. Inferred demographic parameters in dadi using empirical GATK and ANGSD AFS. We analyzed GATK empirical spectra without (dadi) and with low-pass correction (low-pass).

For the isolation-with-migration model fit to YRI and CEU data, the results were broadly similar (Table S1.) For population sizes and the divergence time, inferences were more stable from GATK genotyping and our low-pass model than from ANGSD-estimated AFS. By contrast, the inferred migration rate was similar across analyses.

## Discussion

We assessed the biases introduced by low-pass sequencing using GATK multi-sample genotype calling and developed a model to mitigate these biases. In a simulated population undergoing growth, we found that low-pass sequencing reduced the presence of low-frequency alleles (Fig. 1). Our model accounted for these biases, contrasting with ANGSD, which created fluctuations in the AFS at low depth (Fig. 2). In scenarios involving two populations, we observed similar biases, which our model effectively corrected, whereas ANGSD introduced additional noise (Fig. S3 and S4). For demographic inference, using our model enabled accurate parameter estimates even at low-pass depths, while neglecting low-depth biases resulted in substantial inaccuracies (Fig. 3). ANGSD also yielded accurate estimates, but worse likelihoods. Empirical testing using human data from the 1000 Genomes Project showcased the accuracy of our correction method in improving demographic inference from low-pass data, outperforming both uncorrected analysis and ANGSD results (Fig. 4 and Tables 1 and S1).

While ANGSD is recognized for its effectiveness in managing low-pass sequencing, our results showed its difficulties in modeling medium-frequency alleles. This is reflected in lower likelihood scores, particularly when comparing low-pass datasets to high-pass ones (Fig. 3). Despite their utility in incorporating uncertainty related to low-pass sequencing (Nielsen et al. 2011; Fumagalli 2013; Korneliussen et al. 2014), genotype likelihoods might not always accurately capture the entire range of allele frequencies. Despite the AFS fluctuations, ANGSD yielded reliable parameter estimates for simulated data. But ANGSD was unable to accurately estimate the demographic parameters of real datasets, as demonstrated in the analysis of the 1000 Genomes Project data (Tables 1 and S1). This underscores the need for rigorous and critical assessments of results by evaluating the likelihood of the model and conducting uncertainty analysis.

Variant discovery using GATK involves two main approaches: multi-sample (classic joint-calling) and single-sample calling (Nielsen et al. 2011). We modeled multi-sample calling, which has higher statistical power compared to single-sample calling (Nielsen et al. 2011; Poplin et al. 2018). But multi-sample calling can become computationally burdensome with larger sample sizes, leading to the development of incremental single-calling as a scalable alternative (McKenna et al. 2010; Auwera & O’Connor 2020). When our model was applied to incremental single-calling AFS from subsampled 1000 Genomes Project data, parameter inference was poor (Table S2). Therefore, our model should only be used with multi-sample calling, and a slightly different model may need to be developed for incremental single-calling.

We present a GATK multi-sample calling model designed to compensate for AFS biases introduced by low-pass sequencing. Although tailored for GATK, our model’s design allows for its extension to different pipelines with modifications to address the unique aspects of each calling algorithm. For example, our model currently assumes that a site is called when at least two reads supporting the alternative allele are found (Eq. 3 and 4), but this could be modified for other pipelines with different calling criteria. Our approach can thus be generalized to other calling pipelines, including those using short reads, long reads, and hybrid approaches (e.g. Bankevich et al. 2012; Poplin et al. 2018). Note that our mathematical model assumes a shared read depth distribution among all individuals, and some studies may vary depth among individuals. Simulations suggest, however, that our model remain accurate with uneven depths (Fig. S10).

Our approach can also be integrated into other AFS-based inference tools such as moments (Portik et al. 2017; Leaché et al. 2019), fastsimcoal2 (Excoffier et al. 2013, 2021), GADMA (Noskova et al. 2020), and delimitR (Smith & Carstens 2020), because our approach modifies the model AFS, independent of how it is computed. Our approach may also be useful in Approximate Bayesian Computation (Beaumont 2010; Csilléry et al. 2012) and machine learning workflows (Pudlo et al. 2016; Smith & Carstens 2020), facilitating simulation of low-pass datasets. Note, however, that we model bias in the mean shape of the AFS under low-pass sequencing, not its full variance (Fig. S11). Furthermore, AFS-based analyses are used not only for demographic studies but also to examine natural selection, including inferring the distribution of fitness effects of new mutations (Eyre-Walker & Keightley 2007; Huang et al. 2021). Our approach can thus facilitate population genomics research across tools, approaches, and problem domains.

In conclusion, we have developed a robust correction for low-pass sequencing biases, significantly enhancing the accuracy of demographic parameter estimation at various coverage depths. As the genetic research community continues to address challenges associated with low-pass data (Bryc et al. 2013; Korneliussen et al. 2014; Blischak et al. 2018; Meisner & Albrechtsen 2018), especially when constrained by economics or sample availability, our methodology provides enables more reliable genetic analysis.

## Material and Methods

### Simulating AFS under low-pass sequencing

We used msprime (Kelleher et al. 2016; Baumdicker et al. 2022) to generate SNP datasets via coalescent simulations. We simulated two demographic models. The demographic models were visualized using demesdraw (Gower et al. 2022). The first model, singe-population exponential growth (Fig. S2A), involved two parameters: the relative population size *ν*_1_ = 10 and time of past growth *T* = 0.1 (in units of two times the effect population size generations). The second model, two-population isolation (Fig. S2B), involved three parameters: equal relative sizes of populations 1 and 2, *ν*_1_ = *ν*_2_ = 1, and divergence time in the past T = 0.1. For each model, we conducted 25 independent simulations. For the exponential growth model, we sampled 20 diploid individuals, whereas for the isolation model, we sampled 10 individuals per population. Both demographic scenarios used an ancestral effective population size *N*_e_ of 10,000, a sequence length of 10^7^ bp, a mutation rate of *µ* = 10^−7^ per site per generation, and recombination rate of *r* = 10^−7^ per site per generation.

For simulations incorporating inbreeding, we used SLiM 4 (Messer 2013; Haller & Messer 2023). Datasets were generated under a bottleneck and growth model (Fig. S2C), with a population bottleneck of *ν*_*B*_ = 0.25, followed by a population expansion to *ν*_*F*_ = 1.0. The time of the past bottleneck was set at *T* = 0.2, and the level of inbreeding was varied with *F* ∈ {0.1, 0.5, 0.9}. Inbreeding was introduced using the selfing rate, set to 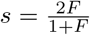. Twenty-five independent simulations were conducted, with 20 individuals sampled for each replicate. Simulation parameters were *N*_*e*_ = 1000, *L* = 2 × 10^6^ bp, *µ* = 5 × 10^−6^, and *r* = 2.5 × 10^−6^, with a burn-in of 10,000 generations.

To create low-pass datasets, we used synthetic diploid genomes. For each simulation replicate, we generated a random reference genome spanning 10 Mb with a GC content of 40%, resembling the human genome. Mutations were incorporated by altering single nucleotides at the positions observed in the SNP matrix generated during each simulation, assuming that all sites were biallelic. Diploid individual genomes were generated by randomly selecting two chromosomes from the population pool.

Using the synthetic individual genomes as templates, we simulated 126 bp paired-end short reads for each individual with InSilicoSeq v2.0.1. (Gourlé et al. 2019). We calculated the number of reads per scenario as *LC/R*, where *L* is the genome length, *C* the coverage depth, and *R* the read length. Reads for each diploid chromosome were simulated with equal probability. Depth of coverage per individual was sampled from a normal distribution with means of 3, 5, 10, and 30 and corresponding standard deviations of 0.3, 0.5, 1, and 3 to explore coverage variability, which increased with coverage levels. These standard deviations were selected based on preliminary simulations that suggested they offer a realistic variance for each coverage level.

For each individual we aligned simulated reads to the reference genome using BWA v0.7.17 (Li et al. 2009). We then processed the aligned reads with SAMTools v1.10 (Li 2013) to perform sorting, indexing, and pileup generation. To generate GATK spectra, we used the GATK multi-sample approach via HaplotypeCaller v4.2 (McKenna et al. 2010; Auwera & O’Connor 2020). To minimize false positives, the identified variants underwent filtering based on GATK’s Best Practices guidelines, with thresholds tailored to expected error rates and variant quality. These thresholds included depth-normalized variant confidence (*QD <* 2.0), mapping quality (*MQ <* 40), strand bias estimate (*FS >* 60.0), and strand bias (*SOR >* 10.0). The filtered SNP VCF files were subsequently used in demographic inference analyses to estimate population parameters based on the AFS of these variants. To generate ANGSD spectra, we used the BAM files containing information about each individual with reads aligned to the reference genome. Subsequently, realAFS was used to estimate a maximum-likelihood AFS through the Expectation-Maximization algorithm. ANGSD v0.94 analysis was executed with the following settings: `doSaf = 1`, `minMapQ = 1`, `minQ = 20`, and `GL = 2`.

### Empirical subsampling of Human data

We used high-quality whole-genome sequencing data (30×) from the 1000 Genomes Project (1kGP), sourced from The International Genome Sample Resource data portal (https://www.internationalgenome.org/ Fairley et al. 2020). The data comprised CRAM files aligned to the GRCh38 human reference genome. We focused on two sets of samples for our analysis: 10 randomly selected individuals from the Yoruba from Ibadan, Nigeria (YRI) samples and 10 from the Utah residents with Northern and Western European ancestry (CEU) samples. The specific individuals included for the YRI were NA18486, NA18499, NA18510, NA18853, NA18858, NA18867, NA18878, NA18909, NA18917, NA18924, and for the CEU NA07037, NA11829, NA11892, NA11918, NA11932, NA11994, NA12004, NA12144, NA12249, NA12273. Additionally, for a single-population demographic model, 20 YRI individuals were analyzed, which includes the initial 10 plus an additional 10 samples: NA19092, NA19116, NA19117, NA19121, NA19138, NA19159, NA19171, NA19184, NA19204, and NA19223.

Initially, we converted the CRAM files to BAM format and indexed them using Picard tools (https://broadinstitute.github.io/picard/). We then isolated reads from chromosome 20 at the original 30 × coverage, which we subsequently subsampled to 10 ×, 5 ×, and 3× coverage using samtools v.1.10 (Li 2013) to emulate varying sequencing depths. Next, using GATK version 4.2.5 HaplotypeCaller (McKenna et al. 2010; Auwera & O’Connor 2020), we called SNPs and indels from these varying coverage depths for each population. We employed multi-sample SNP calling, merging BAM files with identical coverage prior to processing with HaplotypeCaller. This approach yielded a raw output VCF file.

We also carried out a single-sample calling procedure. For this, individual BAM files were used directly as inputs for the GATK HaplotypeCaller with the `-ERC GVCF` flag to enable GVCF mode. Following this, we used GATK GenomicsDBImport to compile the individual variant calls into a cohesive data structure. This setup allowed us to conduct joint genotyping using GATK GenotypeGVCFs, ultimately producing a multi-sample VCF.

Following SNP calling, we employed GATK SelectVariants to filter out indels for both approaches, retaining only SNPs. Quality filtering of SNPs was conducted using GATK VariantFiltration, applying criteria such as depth-normalized variant confidence (*QD <* 2.0), mapping quality (*MQ <* 40), strand bias estimate (*FS >* 60.0), and overall strand bias (*SOR >* 10.0). After quality filtering, the VCF files were annotated with ancestral allele information using the `vcftools fill-aa` module, based on data from the Ensembl Release 110 Database (Danecek et al. 2011).

Finally, we used ANGSD to generate an AFS by using BAM files as input. The sample allele frequencies were first estimated using ANGSD’s `-doSaf` flag, using GATK genotype likelihoods. These likelihoods were then used to calculate the AFS via the Expectation-Maximization algorithm using ANGSD’s realAFS program. In this way, we maintained the original sample sizes from the BAM files, resulting in AFS for 40 chromosomes in the single-population analysis and 20 chromosomes per population in the two-population analysis.

### Demographic inference using dadi

We used dadi (Gutenkunst et al. 2009) to fit demographic models to simulated and empirical datasets. For the GATK spectra, we used the VCF files as input and subsampled individuals to accommodate missing data. For the ANGSD spectra, we used them as input directly. Within dadi, we used three demographic models for the simulated datasets: (i) an exponential growth model: `dadi.Demographics1D.growth`; (ii) a divergence model with migration fixed to zero: `dadi.Demographics2D.split mig`; (iii) an bottleneck then exponential growth model modified to incorporate inbreeding: `dadi.Demographics1D.bottlegrowth`. For the human datasets, we used two models: (i) a divergence with migration model: `dadi.Demographics2D.split mig` and (ii) an instantaneous growth model: `dadi.Demographics1D.two epoch`. The extrapolation grid points were set using the formula [*max*(*ns*) + 120, *max*(*ns*) + 130, *max*(*ns*) + 140], where *ns* is the sample size of the AFS. Our low-coverage correction is also implemented in dadi-cli (Huang et al. 2023).

## Acknowledgements

This work was supported by the National Institute of General Medical Sciences of the National Institutes of Health (R01GM127348 and R35GM149235 to R.N.G.).

## Supplementary Material

**Table S1:** Two-population model analysis results. Inferred demographic parameters in dadi using empirical GATK and ANGSD AFS. We analyzed GATK empirical spectra without (dadi) and with low-pass correction (low-pass).

| parameter | AFS | model | depth |  |  |  |
| --- | --- | --- | --- | --- | --- | --- |
|  |  |  | 30× | 10× | 5× | 3 × |
| $\nu_{YRI}$ | GATK | dadi | 1.79 | 1.63 | 1.18 | 0.61 |
|  | GATK | low-pass | 1.82 | 1.62 | 1.67 | 1.69 |
|  | ANGSD | dadi | 1.69 | 1.58 | 1.26 | 0.87 |
| $\nu_{CEU}$ | GATK | dadi | 0.38 | 0.37 | 0.31 | 0.17 |
|  | GATK | low-pass | 0.38 | 0.38 | 0.36 | 0.34 |
|  | ANGSD | dadi | 0.39 | 0.38 | 0.33 | 0.22 |
| $T$ | GATK | dadi | 0.21 | 0.22 | 0.18 | 0.06 |
|  | GATK | low-pass | 0.21 | 0.23 | 0.20 | 0.16 |
|  | ANGSD | dadi | 0.21 | 0.22 | 0.20 | 0.07 |
| $m$ | GATK | dadi | 1.80 | 2.00 | 2.24 | 1.68 |
|  | GATK | low-pass | 1.80 | 2.00 | 1.89 | 1.66 |
|  | ANGSD | dadi | 1.99 | 2.12 | 2.44 | 1.91 |
| $\theta$ ($\times 10^4$) | GATK | dadi | 5.42 | 5.44 | 5.56 | 5.65 |
|  | GATK | low-pass | 5.43 | 5.42 | 5.45 | 5.40 |
|  | ANGSD | dadi | 6.04 | 6.01 | 6.03 | 6.25 |
| log-likelihood | GATK | dadi | -2588 | -2378 | -2329 | -2663 |
|  | GATK | low-pass | -2590 | -2479 | -2224 | -1850 |
|  | ANGSD | dadi | -5518 | -5595 | -7074 | -11029 |

**Table S2:** One-population model analysis results with single-sample calling using empirical GATK AFS. We analyzed GATK empirical single-sample call spectra without (dadi) and with low-pass correction (low-pass).

| parameter | model | depth |  |  |  |
| --- | --- | --- | --- | --- | --- |
|  |  | 30× | 10× | 5× | 3 × |
| $\nu_{YRI}$ | dadi | 1.85 | 1.87 | 1.82 | 1.56 |
|  | low-pass | 1.86 | 1.93 | 2.73 | 3.60 |
| $T$ | dadi | 0.43 | 0.45 | 0.51 | 0.48 |
|  | low-pass | 0.42 | 0.40 | 0.24 | 0.24 |
| $\theta$ ($\times 10^3$) | dadi | 5.13 | 5.05 | 4.62 | 4.31 |
|  | low-pass | 5.14 | 5.10 | 4.96 | 4.49 |
| log-likelihood | dadi | -284 | -280 | -457 | -1755 |
|  | low-pass | -291 | -317 | -597 | -1005 |

**Figure S1:**
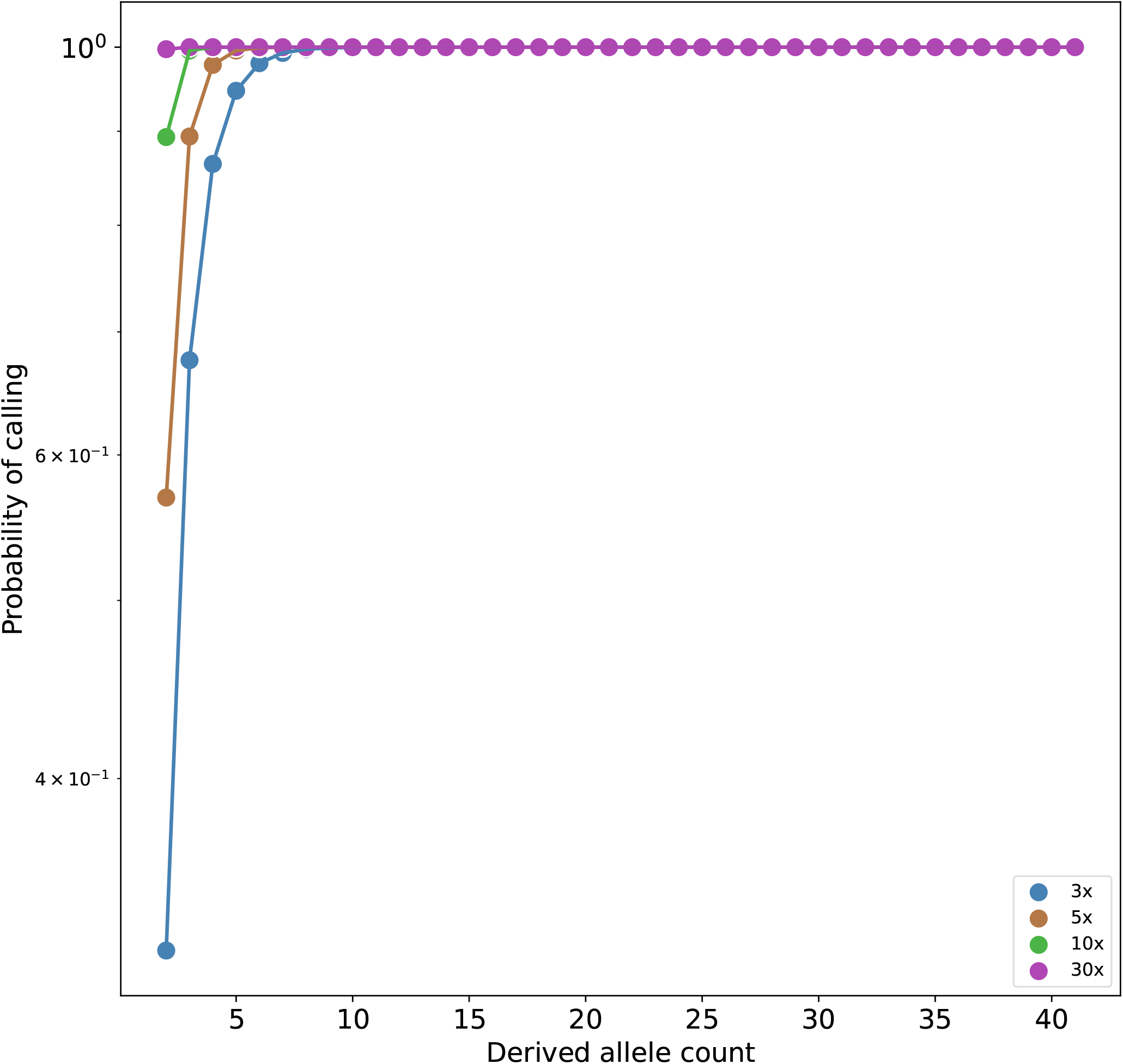
Probability of calling a variant site versus true allele frequency and coverage depth.

**Figure S2:**
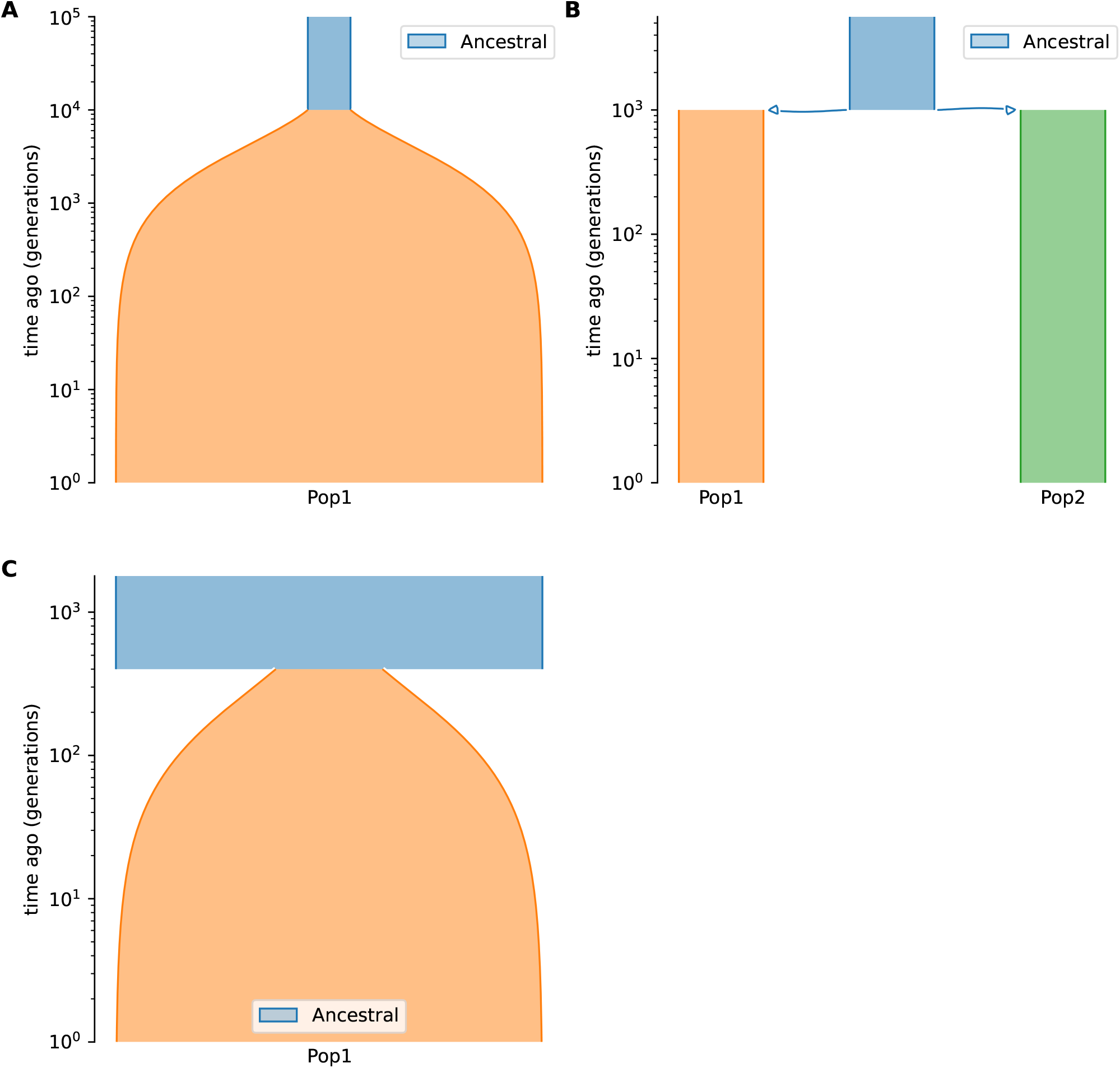
Representation of the demographic models used in the simulations: (A) single-population exponential growth model with parameters *ν*_1_ = 10 and *T* = 0.1, (B) two-population isolation model with *ν*_1_ = *ν*_2_ = 1 and *T* = 0.1, (C) single-population exponential growth model with inbreeding with parameters *ν*_1_ = 4, *T* = 0.4, and *F* ∈ {0.1, 0.5, 0.9}. *ν, T, F* represent relative population size, time in the past, and inbreeding coefficient, respectively. This plot was created with Demes (Gower et al. 2022)

**Figure S3:**
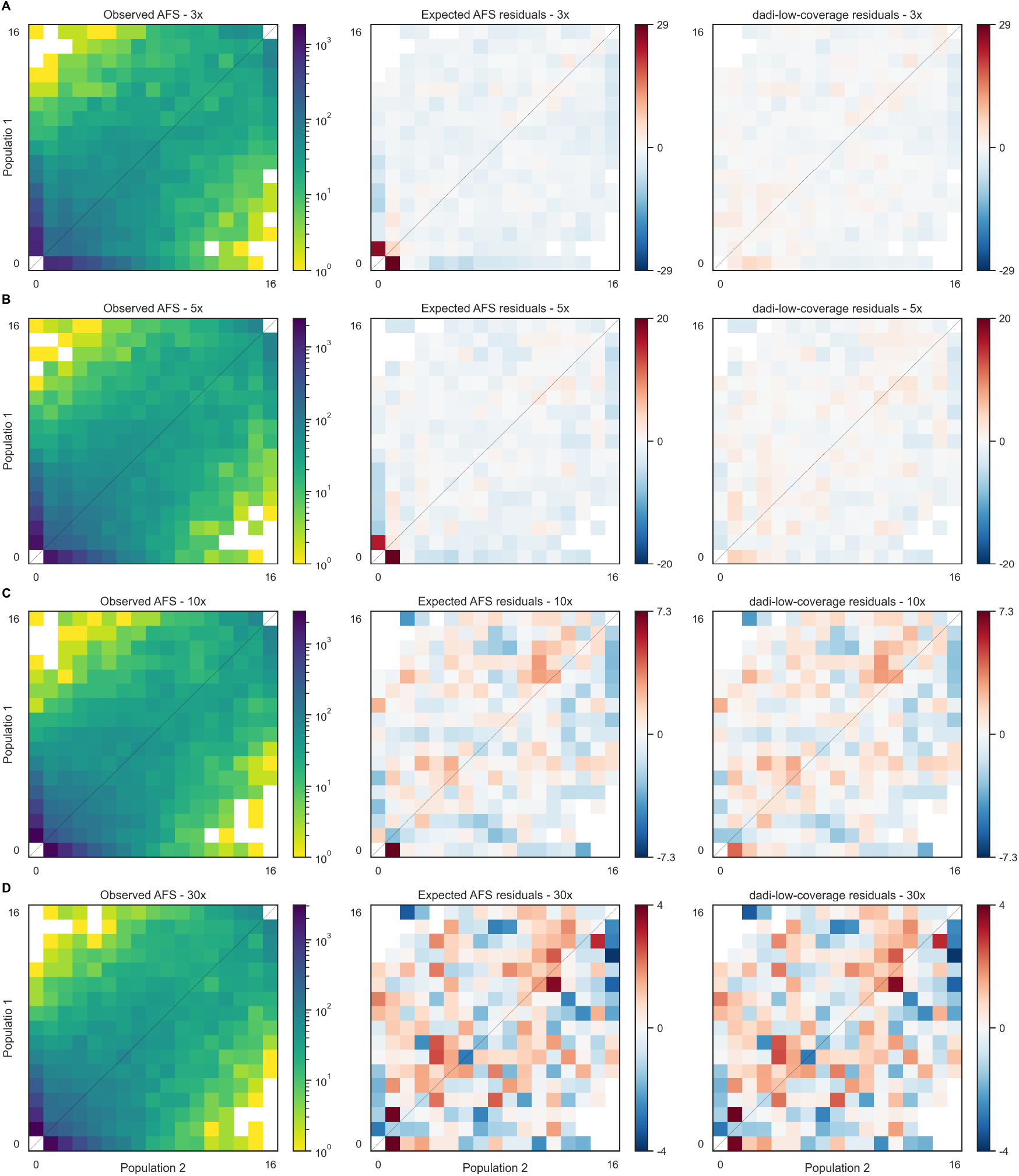
The observed 2D AFS is biased by low coverage. Deviation between the observed low-coverage AFS (first column) and the expected AFS (calculated by dadi) for the isolation demographic scenario is visualized through the residual plot (second column). Dark red residuals indicate that the observed low-coverage AFS is deficient in low-frequency alleles compared to the expectation. By contrast, the residuals between the observed AFS and the low-coverage model are much smaller. At 30× coverage (D) the residuals become small and random, indicating agreement between all three spectra. Coverage depths compared are (A) 3×, (B) 5×, (C) 10×, and (D) 30×.

**Figure S4:**
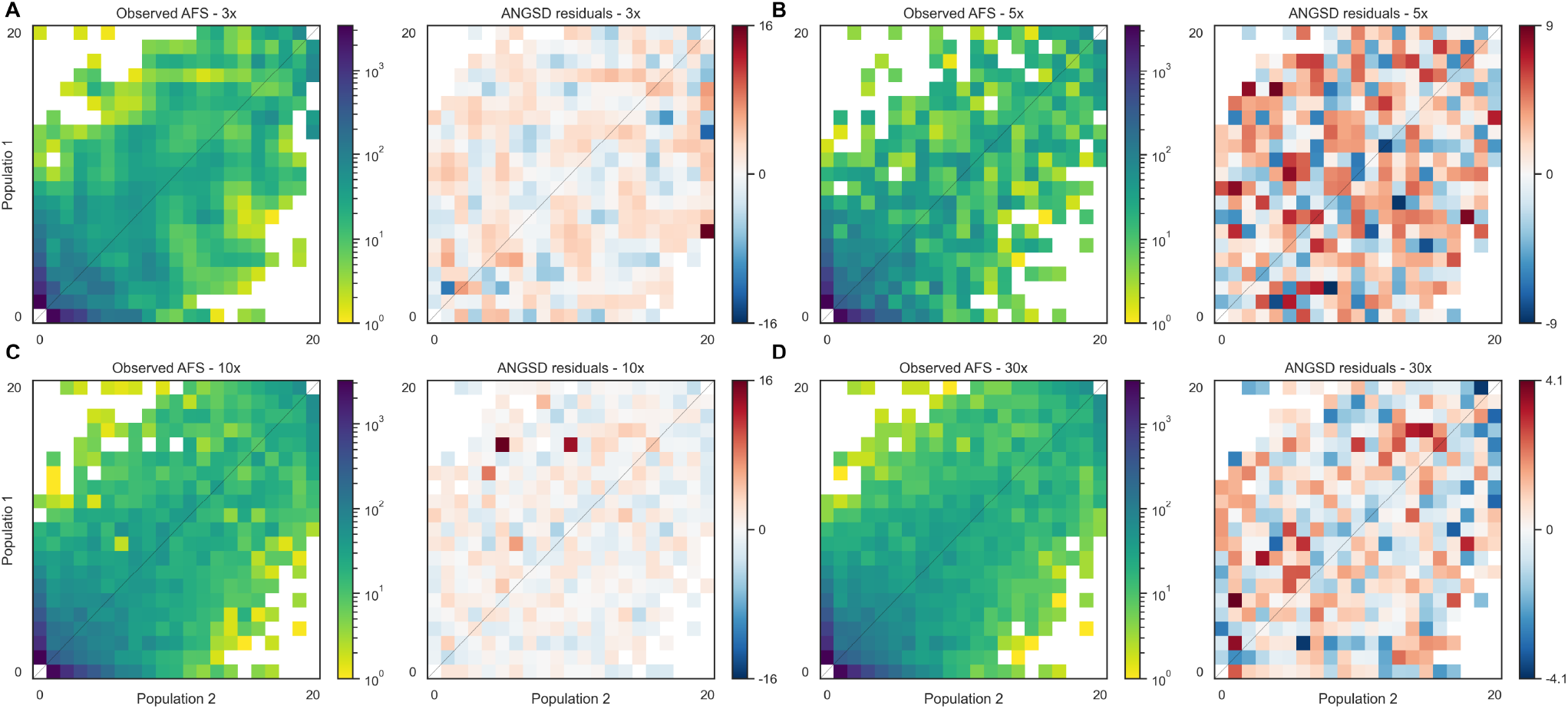
ANGSD creates fluctuations in the joint AFS. The joint AFS output by ANGSD exhibits sporadic very large residuals when compared with the true simulated AFS, similar to the oscillations seen in the single population AFS (Fig. 2). Coverage depths compared are (A) 3×, (B) 5×, (C) 10×, and (D) 30×.

**Figure S5:**
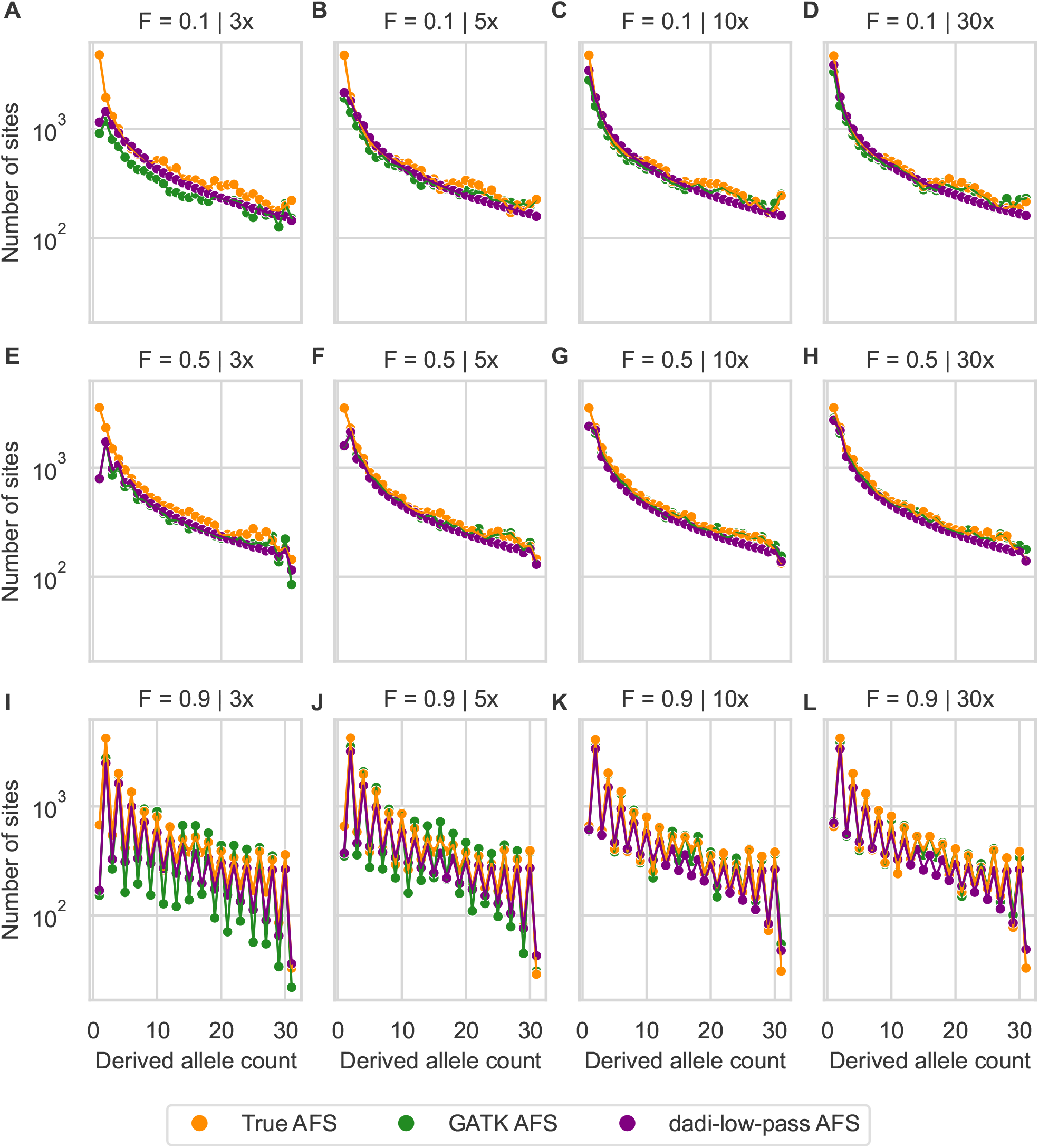
The observed AFS is impacted by low-pass sequencing (3×, 5×, 10×, and 30×) and inbreeding (*F* ∈ {0.1, 0.5, 0.9}). This figure presents a comparison of the observed AFS from low-pass variant calling with simulations in both the standard dadi and dadi-low-pass frameworks, using the true parameter values for a single-population model.

**Figure S6:**
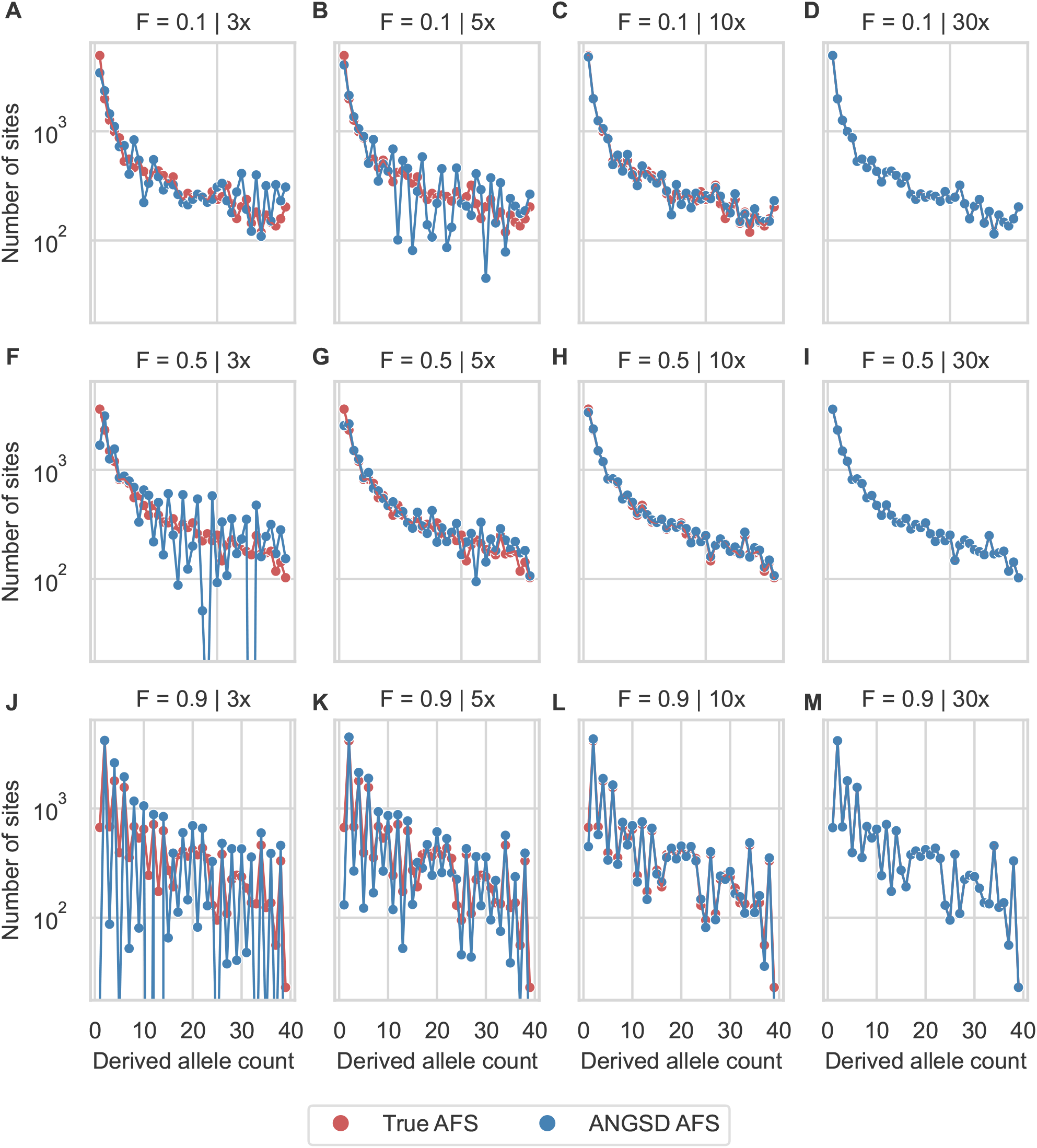
ANGSD corrects for the low-pass bias of the AFS, but it introduces fluctuations in inbreeding models. For the same simulations as Fig. S5, ANGSD (blue) was used to reconstruct the simulated AFS (red). Coverages were 3×, 5×, 10×, and 30×) and inbreeding 0.1, 0.5, and 0.9.

**Figure S7:**
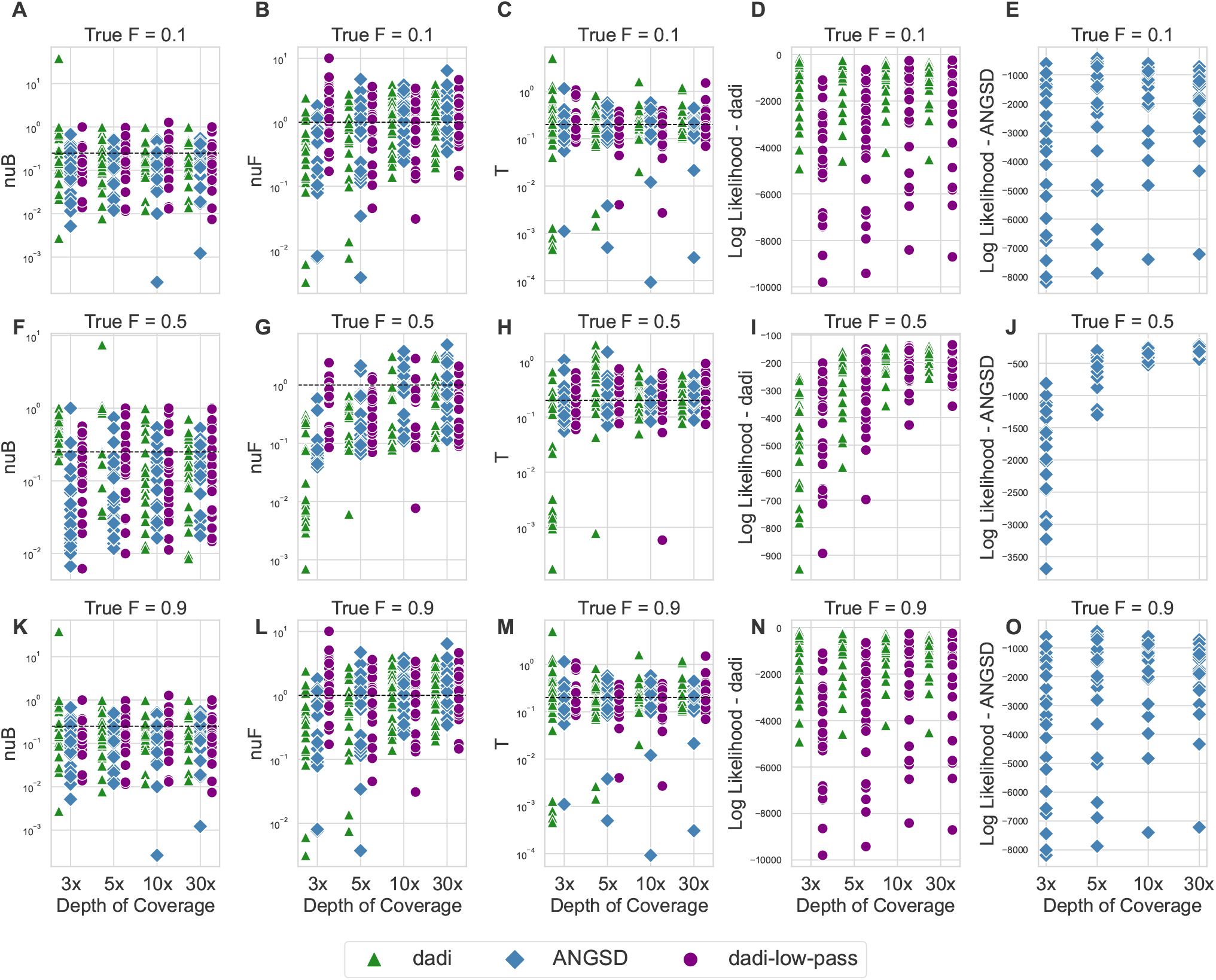
Graph showcasing the accuracy of parameter and likelihood estimations across various sequencing depths (3×, 5×, 10×, and 30×) and inbreeding (*F* ∈ {0.1, 0.5, 0.9}) for a population bottleneck and growth model. The inbreeding parameters were kept fixed for both the low-pass calculation and the optimization process. Parameters were obtained through different methods, including dadi, both with and without corrections for low coverage, as well as ANGSD. Details of the graph include: (A), (F), (K) the estimated size after population bottleneck; (B), (G), (L) the estimated size after population expansion; (C), (H), (M) the time of population expansion; (D), (I), (N) log-likelihood calculations from dadi, highlighting the distinction between corrected and uncorrected model for low coverage; and (E), (J), (O) log-likelihood calculations from ANGSD. The black line present in the plots for (A), (B), (E), (F), (I), (J) and indicates the true value of the parameter, providing a standard for evaluating the accuracy of different approaches.

**Figure S8:**
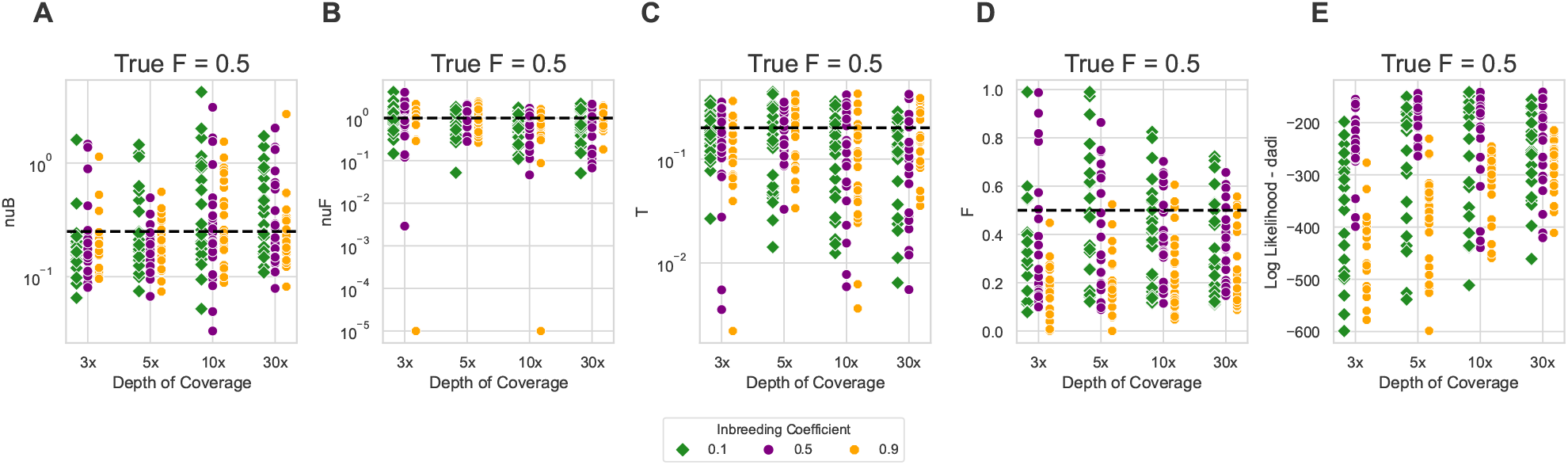
Graph showcasing the accuracy of parameter and likelihood estimations across various sequencing depths (3 ×, 5 ×, 10×, and 30×) and inbreeding (∈{*F* 0.1, 0.5, 0.9}) for a population expansion model under a true inbreeding value of 0.5. The inbreeding parameters used for the low-pass calculation were 0.1, 0.5, and 0.9. Parameters were obtained using dadi-low-pass. Details of the graph include: (A) the estimated size after population bottleneck; (B) the estimated size after population expansion; (C) the time of population expansion; (D) inferred inbreeding coefficient; (E) log-likelihood calculations from dadi-low-pass. The black line present in the plots for (A), (B), (C), and (D) indicates the true value of the parameter, providing a standard for evaluating the accuracy of different approaches.

**Figure S9:**
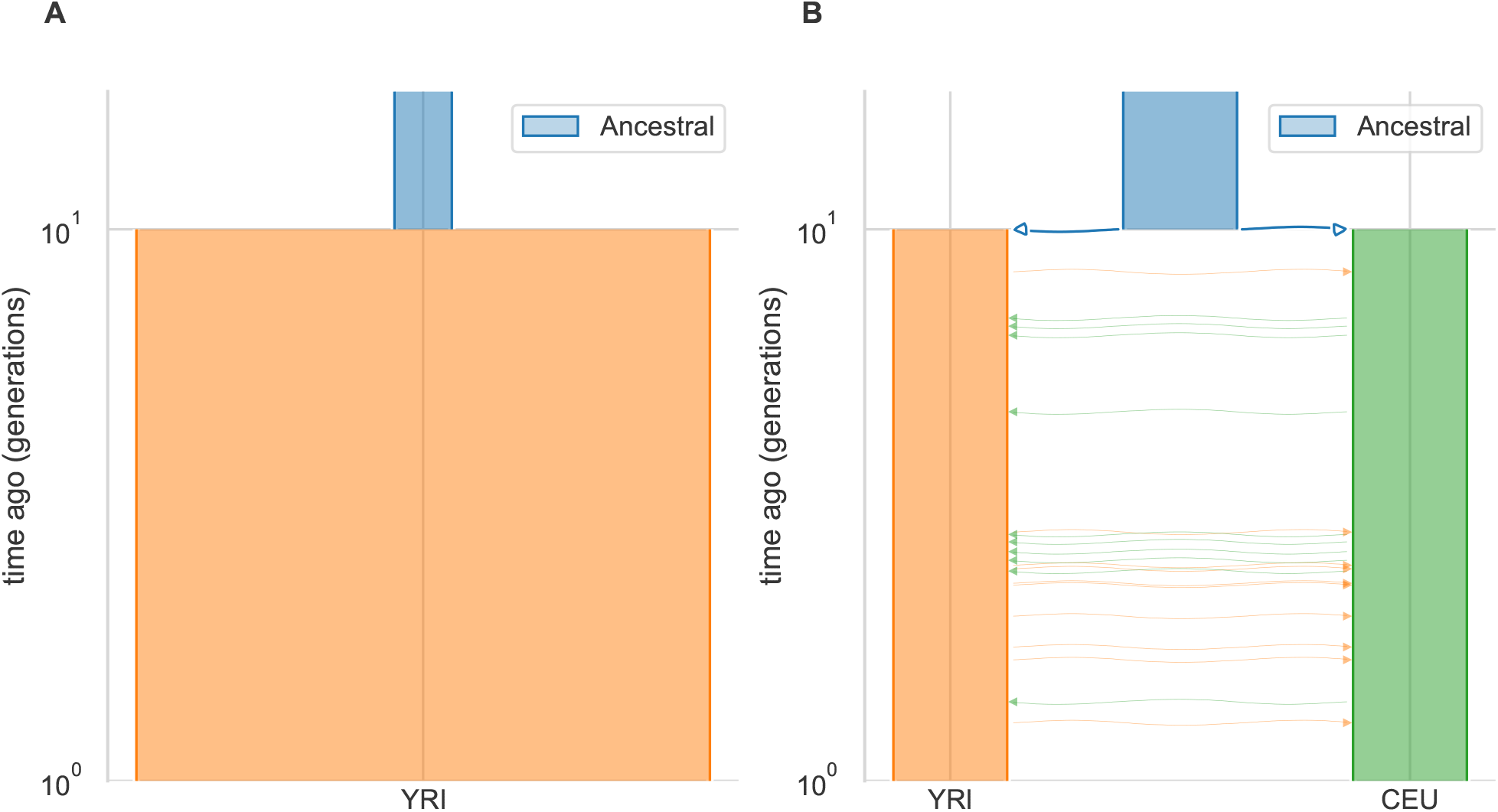
Representation of the demographic models used to analyse 1000 genomes datasets: (A) single-population two-epoch growth model with parameters, (B) two-population isolation with migration model. This plot was created with Demes (Gower et al. 2022)

**Figure S10:**
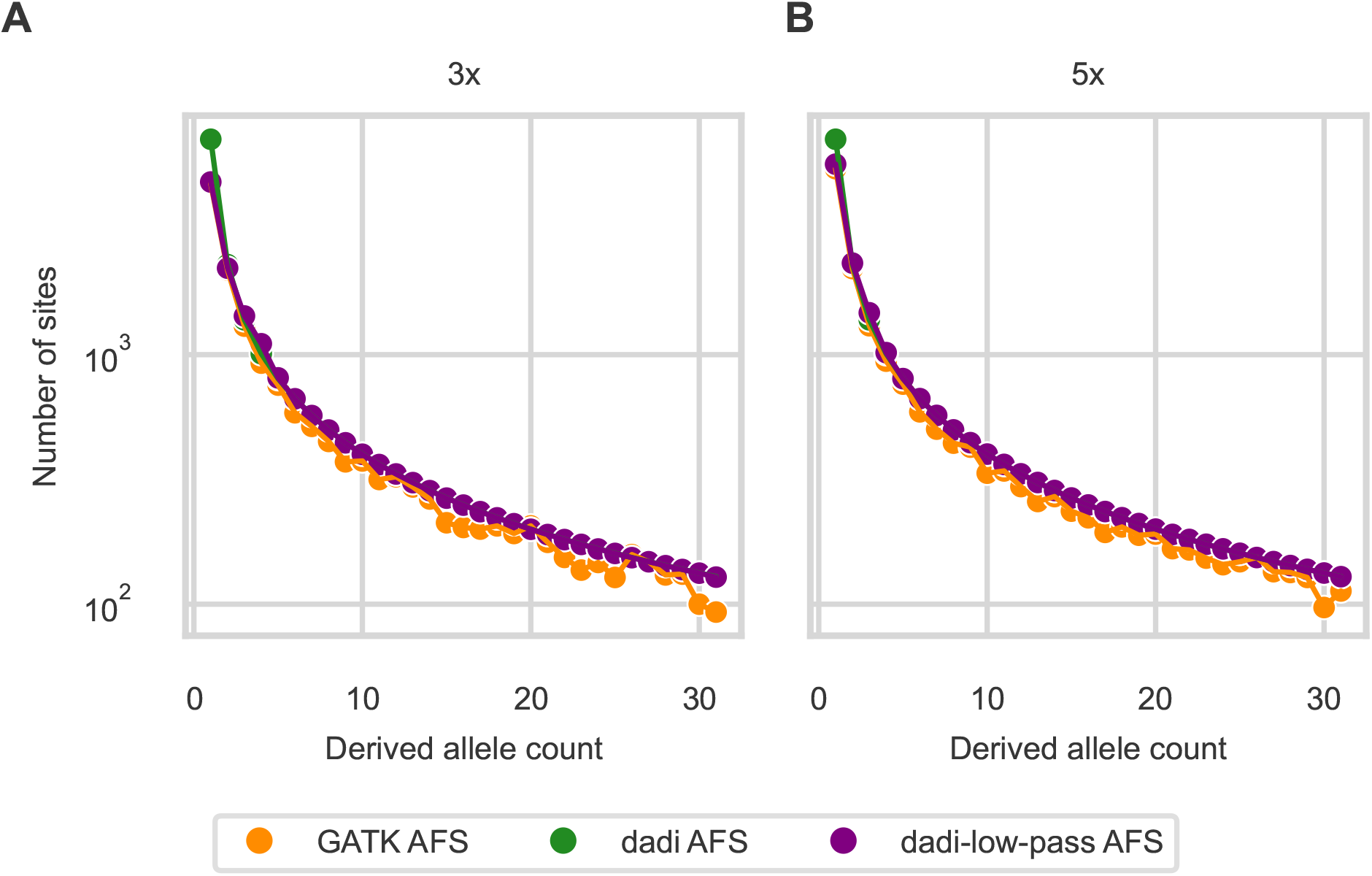
Unbalanced depth of coverage does not bias the dadi-low-pass model. Simulations were performed using 20 individuals, with half simulated under low-coverage conditions (A: 3× or B: 5×) and the other half under high-depth coverage (30 ×).

**Figure S11:**
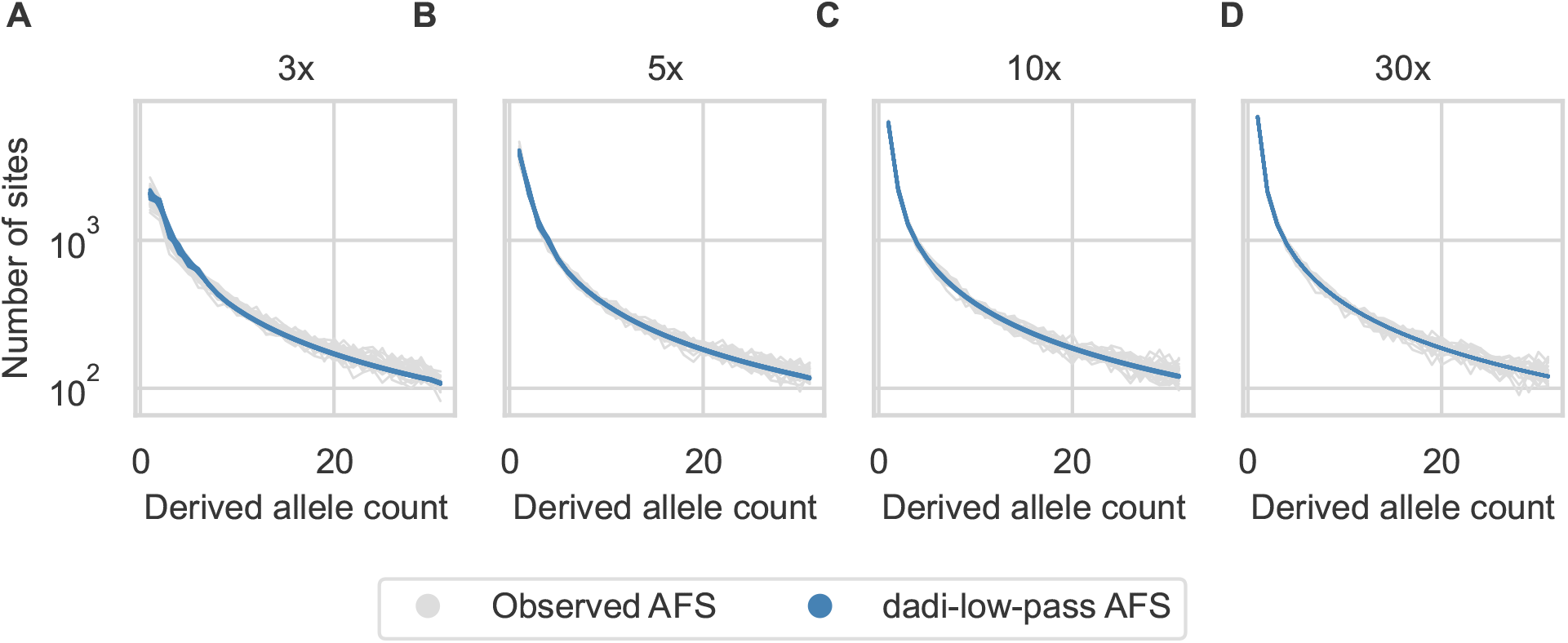
The simulated AFS under the low-pass model shows less variance compared to that observed in the simulated datasets. We generated 25 AFS for each condition.

